# Hygrochastic Seedpods: Engineering Microclimates for Forest Restoration

**DOI:** 10.64898/2026.09.12.747887

**Authors:** Semina Yi, Yaoye Hong, Florentin C. Jaeger, Marco Freschi, Yaobin Yang, Ziyun Zhang, Hajun Lee, Kenichi Soga, Ehren Moler, Teng Zhang, Taryn L. Bauerle, Shu Yang, Lining Yao

## Abstract

Forest restoration at scale depends on sowing seeds directly onto the land, yet seedling establishment is often poor and inconsistent. Seed-enhancement technologies — priming, coating, and pelleting — improve germination under water, temperature, and salinity stress by supplying polymers, nutrients, or microorganisms at the seed surface. Being mainly conformal films, they act continuously once wetted and cannot gate when a seed is exposed to the soil. They also enclose no air volume, so the microenvironment they create is inseparable from the surrounding soil and has been inferred from plant performance rather than measured directly. Whether an engineered carrier can instead set both the state and the duration of the post-dispersal microenvironment, and how such an effect can be directly quantified, remains unexplored. Here we show that hygrochastic seedpods hold seeds in a timed, directly measurable microclimate and accelerate seedling emergence. The pods are programmable, responsive conical shells enclosing an air cavity around the seed. Two design solutions were investigated in parallel: wooden seedpods that crack open upon hydration, and poly(vinyl alcohol)/boric acid seedpods that gradually disintegrate after sustained wetting. In both, wax coatings act as a programmable diffusion-limited timer that sets activation from hours to days. The pod cavity permits direct sensing and monitoring, which no coating geometries allow. As directly measured during the initial phase after deployment, seedpod interiors ran 2–3 °C warmer and drier than the surrounding soil, and these differences persisted after opening. Across indoor and field trials in several tree species, pods accelerated early seedling emergence without penalizing subsequent growth. These results identify artificial hygrochastic seedpods as a transient ecological mediator that regulates both the timing of seed exposure and the microclimate around the seed, offering a strategy for improving forest restoration under variable environmental conditions.

---

Ecological restoration is essential for sustaining ecosystems, securing natural resources, and preserving biodiversity. Yet restoration success is frequently limited by poor seedling establishment through direct seeding, where only a small fraction of dispersed seeds successfully establishes as mature seedlings^1^. Seedling transplanting improves establishment rates but is time- and budget-intensive, which limits its scalability. Direct seeding therefore remains the competitive practical route to landscape-scale restoration. To improve its yield, seed-enhancement technologies —priming, coating, encrusting, pelleting, and agglomeration —have been adapted from agricultural practice to native and restoration species ^2–4^. These treatments supplement the seed with water-holding polymers, nutrients, or beneficial microorganisms, and improve germination under water, temperature, and salinity stress ^5,6^. These treatments, however, have two limitations. First, a coating acts on the soil conditions immediately available at the seed surface and, once wetted, acts continuously —it does not gate when the seed becomes exposed to the surrounding soil. Second, a conformal thin film cannot enclose a volume of air. The microenvironment it creates is therefore inseparable from the surrounding soil and cannot be adjusted independently of it. It also cannot be measured: with no interior cavity to hold a sensor, its benefit has been inferred from plant performance rather than observed directly^2,6^. For tree seeds in forest restoration, where establishment often depends on a single unpredictable wetting window, both the timing of exposure and the conditions experienced before it may matter. Gating exposure and regulating the enclosed microclimate are the focus of this work.

Natural seed-bearing structures provide models for temporal gating of seed exposure. Among the diverse strategies that synchronize seed dispersal with environmental conditions, hygrochasy provides a striking example that triggers fruits or capsules opening by moisture or water absorption^7^ to release seeds at environmentally optimal times. Specifically, hygrochastic structures remain closed under transient humidity but open only after sustained wetting, synchronizing seed release with rainfall events that favor germination and seedling establishment (Fig. 1a)^8–10^. Beyond regulating release timing, these structures protect seeds from predators, desiccation, and unfavorable environmental fluctuations before germination^11–15^. For example, *Chorizanthe rigida* relies on hygrochastic dispersal, timing its seed release with episodic rainfall in the desert^16^. Similarly, species in the *Mesembryanthemum* and Aizoaceae families open their fruit capsules upon water absorption, releasing seeds directly into moist soil^17^ (Supplementary Note 1). Such post-dispersal regulation has evolved across diverse plants and increases the likelihood that seeds encounter a favorable microsite for successful establishment.

**Figure 1.**
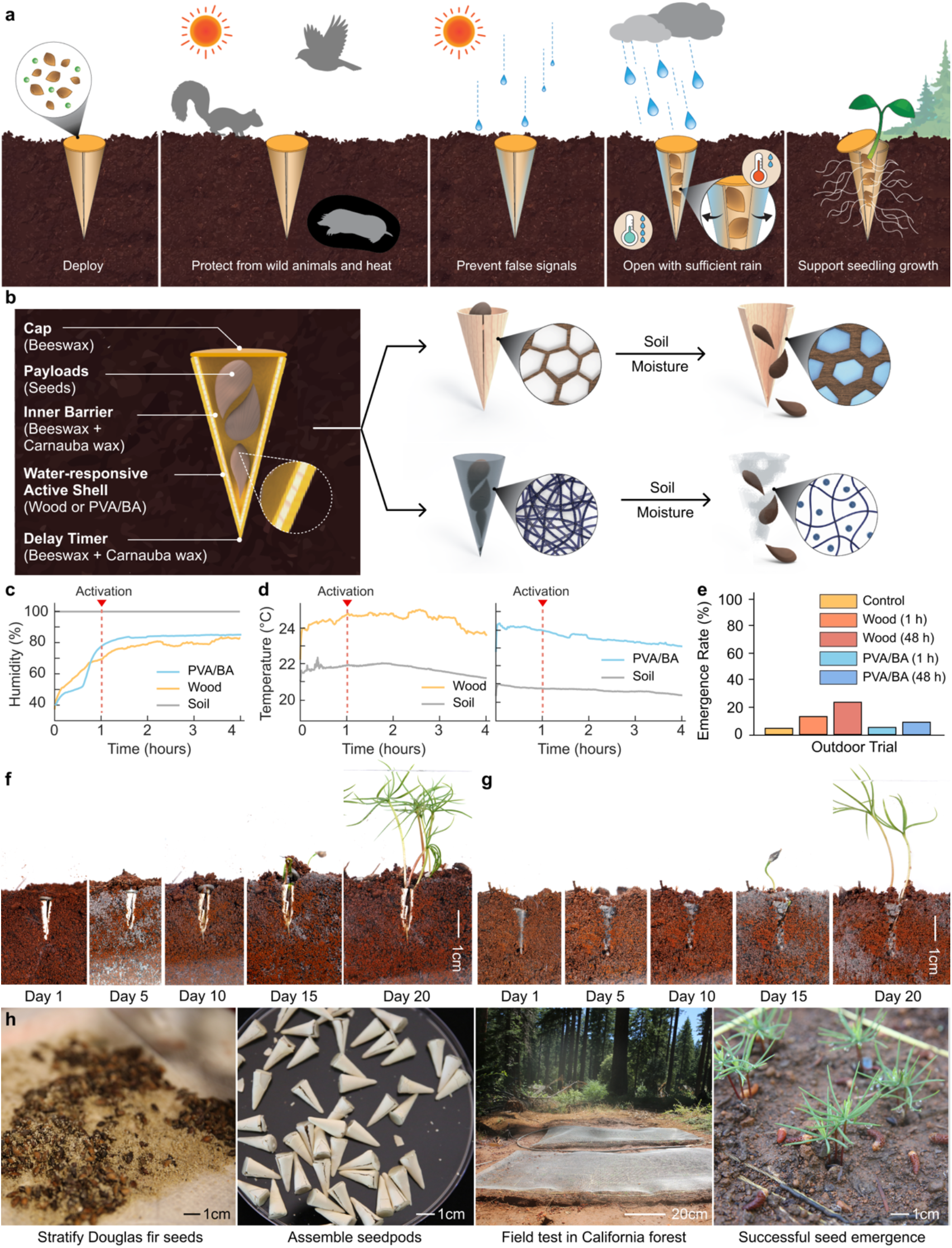
Engineered seedpods regulate the post-dispersal seed microenvironment to promote tree establishment. **a.** Conceptual illustration of the seedpods’ life cycle after deployment. **b.** Cross-sectional view of the conical seedpod design, showing a beeswax cap, seed payloads, an inner beeswax-carnauba wax barrier, a water-responsive active shell, and an outer time-delay wax layer. Soil moisture causes the wooden shell to swell and crack open (top right) or the poly(vinyl alcohol)/boric acid (PVA/BA) shell to dissolve (bottom right), exposing internal pore networks to release the seeds. **c-d.** Representative time series of relative humidity **(c)** and temperature **(d)** inside wooden and PVA/BA seedpods and in the surrounding soil. Red triangles and dashed vertical lines indicate activation at 1 h. **e.** Douglas fir seedling emergence on Day 12 in outdoor tests. **f**-**g**. Time-lapse sequence of the opening mechanism during germination of wooden seedpods (**f**) and PVA/BA seedpods (**g**) in soil under a controlled indoor environment. **h.** Workflow of the full deployment pipeline in the outdoor tests.

Recent engineered systems inspired by natural seeds have improved seed deployment efficiency, landing and soil engagement^18–23^. However, these technologies have largely focused on the physical process of deployment. Whether engineered structures can regulate the post-dispersal microenvironment to promote germination and early establishment remains largely unexplored. Drawing on the temporal gating exhibited by natural hygrochastic structures, we hypothesized that active regulation of the seed microenvironment could improve tree seed germination and early establishment.

To test this hypothesis, we developed two complementary moisture-responsive seedpod strategies: hygromorphic crack-opening wooden seedpods (Supplementary Video 1)^24–26^ and dissolution-open poly(vinyl alcohol)/boric acid (PVA/BA) seedpods that regulate post-dispersal environmental conditions through distinct opening mechanisms (Fig. 1b)^27–29^. Hydrophobic wax coatings delayed activation, allowing the duration of seed enclosure to be tuned while modifying the local humidity (Fig. 1c) and temperature (Fig. 1d) surrounding the seeds. We then investigated how the engineered post-dispersal environments influenced germination, seedling establishment, and early growth through both controlled laboratory experiments and field validation (Fig. 1e-h).

Our studies demonstrate that engineered hygrochastic seedpods successfully regulate the post-dispersal seed microenvironment, with wooden and PVA/BA seedpods maintaining higher temperature and lower relative humidity than the surrounding soil. These microclimate modifications promote earlier seed germination and seedling emergence, with complementary strategies offering flexible options for diverse restoration contexts.

## Result

### Engineered seedpods establish distinct post-dispersal microclimates

As a biological reference, we examined the hygromorphic opening behavior of naturally occurring *Erodium gruinum* seeds, which progressively open over approximately 15 h following hydration (Fig. 2a and Supplementary Video 2). Inspired by this natural strategy, we developed two complementary moisture-responsive seedpod strategies based on distinct activation mechanisms. The wooden seedpod translates natural hygrochastic opening through hydration-induced crack formation, whereas the PVA/BA seedpod gradually dissolves in water following prolonged soil wetting (see Method, Supplementary Note 2 and Supplementary Fig. 1-5). Both seedpods share a common conical geometry and programmable activation enabled by hydrophobic wax coatings (Supplementary Fig. 6), allowing direct comparison of their effects on the post-dispersal environment experienced by seeds.

**Figure 2.**
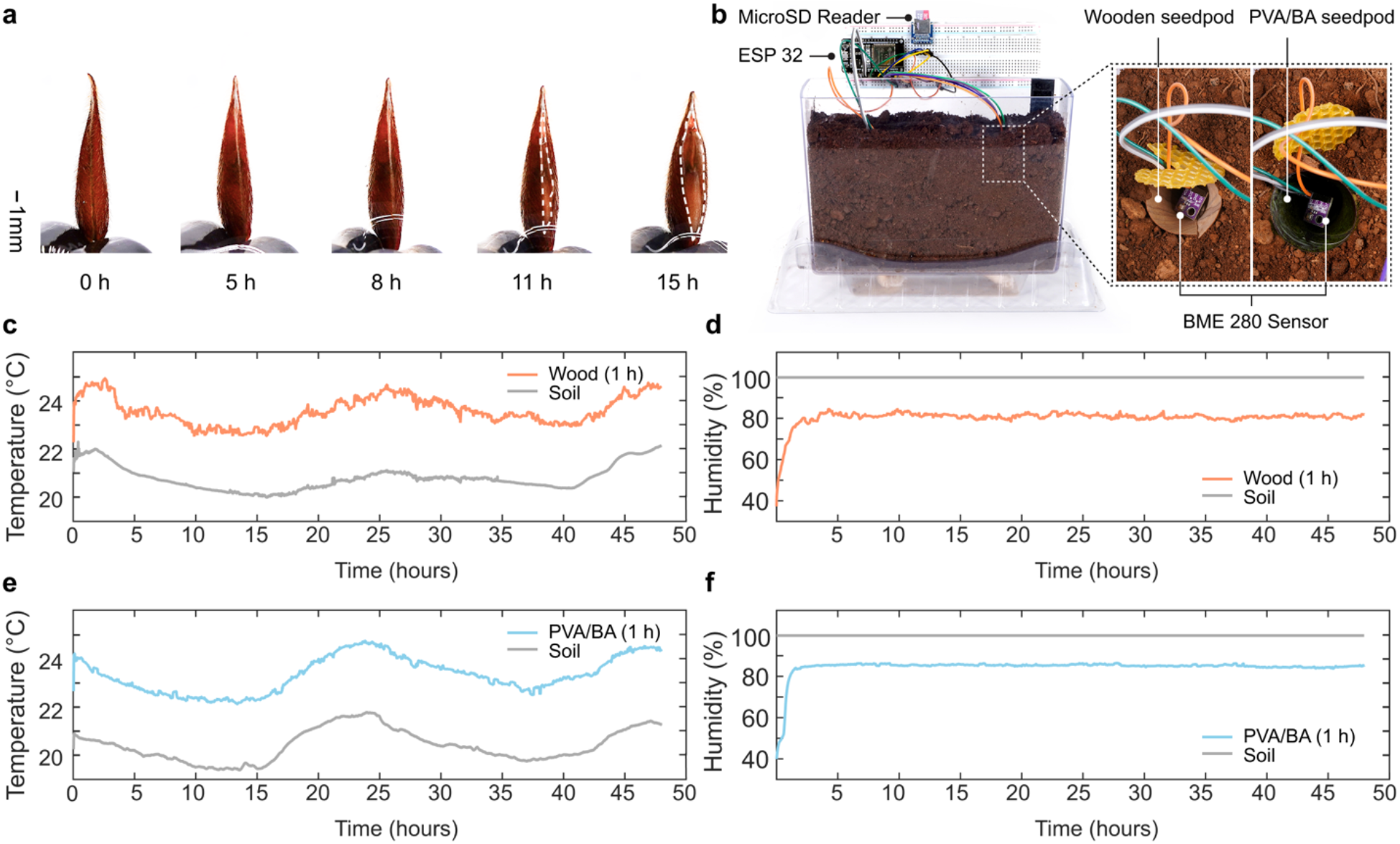
Moisture-responsive seedpods provide complementary strategies for post-dispersal environmental regulation. **a.** Time-lapse of hygromorphic opening in natural *Erodium* seeds, highlighting the progressive opening of the suture over time. **b.** Microclimate chamber used to continuously monitor temperature and relative humidity within the pod and the surrounding soil. **c.** Temperature comparison between soil and the interior of the wooden seedpods, and **d.** relative humidity, measured over a 48-hour period for one representative dataset. **e.** Temperature and **f.** relative humidity measurements for the PVA/BA seedpod show similarly distinct micro-environment profiles relative to the surrounding soil.

The seedpod geometry was adapted from the tapered form of natural *E. gruinum* seeds. with an adjusted internal capable of accommodating up to three tree seeds of target species commonly used in ecological restoration, including Douglas-fir (*Pseudotsuga menziesii*), sweet birch (*Betula lenta*), eastern white pine (*Pinus strobus*), and eastern hemlock (*Tsuga canadensis*). For all our experiments, we fabricated the seedpods to a ∼ 370 mm³ internal volume unless otherwise specified. A predefined longitudinal seam directed controlled opening, whereas the pointed conical geometry facilitated soil penetration and maintained intimate seed–soil contact after deployment. This shared architecture enabled direct comparison of the two moisture-responsive activation strategies while minimizing geometric effects on seed establishment.

For sensor-based microclimate measurements, we used geometrically scaled seedpods with an enclosed volume of 2.59 cm³ to accommodate the sensors (Fig. 2b and Supplementary Note 3). Both seedpod strategies establish microclimates distinct from the surrounding soil. Wooden seedpods maintained temperatures ∼ 2-3°C higher than soil while stabilizing relative humidity (RH) at ∼ 80% in the closed state. Similarly, PVA/BA seedpods maintained elevated temperatures and RH near 85%, whereas RH surrounding the soil is close to saturation. These distinct microclimatic conditions persisted after seedpod opening, with internal temperatures remaining higher and RH remaining lower than those in the surrounding soil, respectively, throughout the monitoring period. Thus, opening exposed the seeds without immediately eliminating the seedpod-associated microclimate.

### Programmable activation enables temporal regulation of the post-dispersal environment

Seedpod opening changes the physical boundary between the enclosed seed and the surrounding soil. Activation timing, therefore, defines how long the engineered microclimate is maintained before seed release (Fig. 2). We next investigated how this duration can be programmed after deployment (Supplementary Figure 8). The wooden seedpod opens through hydration-induced crack propagation, whereas the PVA/BA seedpod gradually disintegrates after prolonged wetting (Fig. 3a,b). Despite the difference in activation mechanisms, both strategies delay seed release through a multilayer architecture consisting of an active shell, an inner moisture barrier, and a hydrophobic wax coating that functions as a programmable delay timer (Fig. 3c).

**Figure 3.**
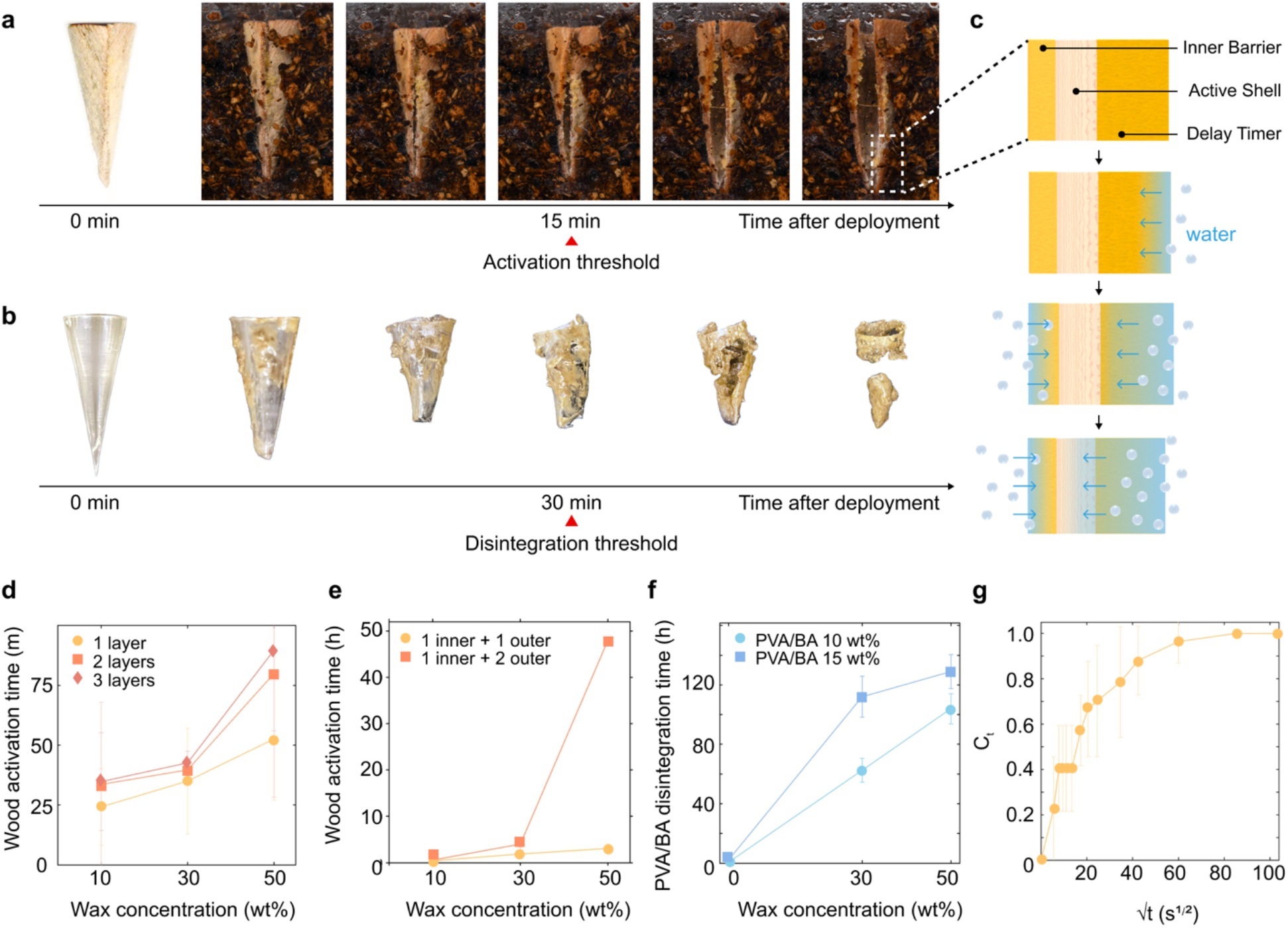
Soil-moisture activation dynamics of the seedpods. **a.** Time-lapse photos of the wooden seedpods in soil, showing moisture uptake, swelling of the active shell, and crack-open activation threshold. **b.** Corresponding time-lapse photos of the PVA/BA seedpods, showing progressive softening, collapse, and eventual dissolution at the activation threshold. **c.** Schematic of the multilayer architecture showing the inner barrier, water-responsive active shell, and outer wax delay-timer, and the corresponding moisture-activation pathway. **d.** Activation time profile of the wooden seedpods with 1–3 layers of a 1:1 mixture of beeswax and carnauba wax coating as a function of wax concentration (n = 5 for each condition, mean ± s.d.). **e.** Activation time profile of the wooden seedpods with combined inner–outer wax configurations as a function of wax concentration (n = 5 for each condition, mean ± s.d.). **f.** Disintegration time of the PVA/BA seedpods as a function of wax concentration (n = 3 for each condition, mean ± s.d.). **g.** Moisture-diffusion analysis of the wax-coated shells to estimate diffusion kinetics across samples (n = 7, mean ± s.d.). Concentration at time t, Cₜ = (Mₜ − M₀) / (M_∞_ − M₀), plotted against the square root of time, √t (s^1/2^).

Increasing wax thickness progressively delayed activation. For wooden seedpods, incorporating an additional inner wax barrier prolonged activation from approximately 3 h to 48 h (Fig. 3d,e). Similarly, increasing wax concentration delayed PVA/BA disintegration from approximately 60 h to 129 h (Fig. 3f).

To examine the delay kinetics, we modeled water transport through the wax coating as a diffusion-limited process. The normalized moisture uptake initially increased approximately linearly with the square root of before approaching saturations, consistent with Fickian diffusion through the hydrophobic barrier (Fig. 3g and Supplementary Note 4). Clearly, wax architecture provided programmable temporal control over how long seeds remain enclosed depending on the thickness and placement of the wax layer; the seeds can be exposed to the surrounding soil from hours to days.

### Engineered post-dispersal environments differentially influence seed germination and establishment

We conducted two consecutive indoor germination tests to assess performance across distinct restoration-relevant species (Supplementary Note 5). One test included four tree species, *Betula lenta, Tsuga canadensis, Acer rubrum, and Pinus strobus*, which are important to U.S. East Coast reforestation efforts. The second test included *Pseudotsuga menziesii* seeds sourced from northern California (Fig. 4a and Supplementary Fig. 11). Both experiments compared wooden and PVA/BA seedpods programmed for 1 h or 48 h activation delay with a direct-sown control. Here, activation delay refers to the duration of wetting before seedpod activation or disintegration.

**Figure 4.**
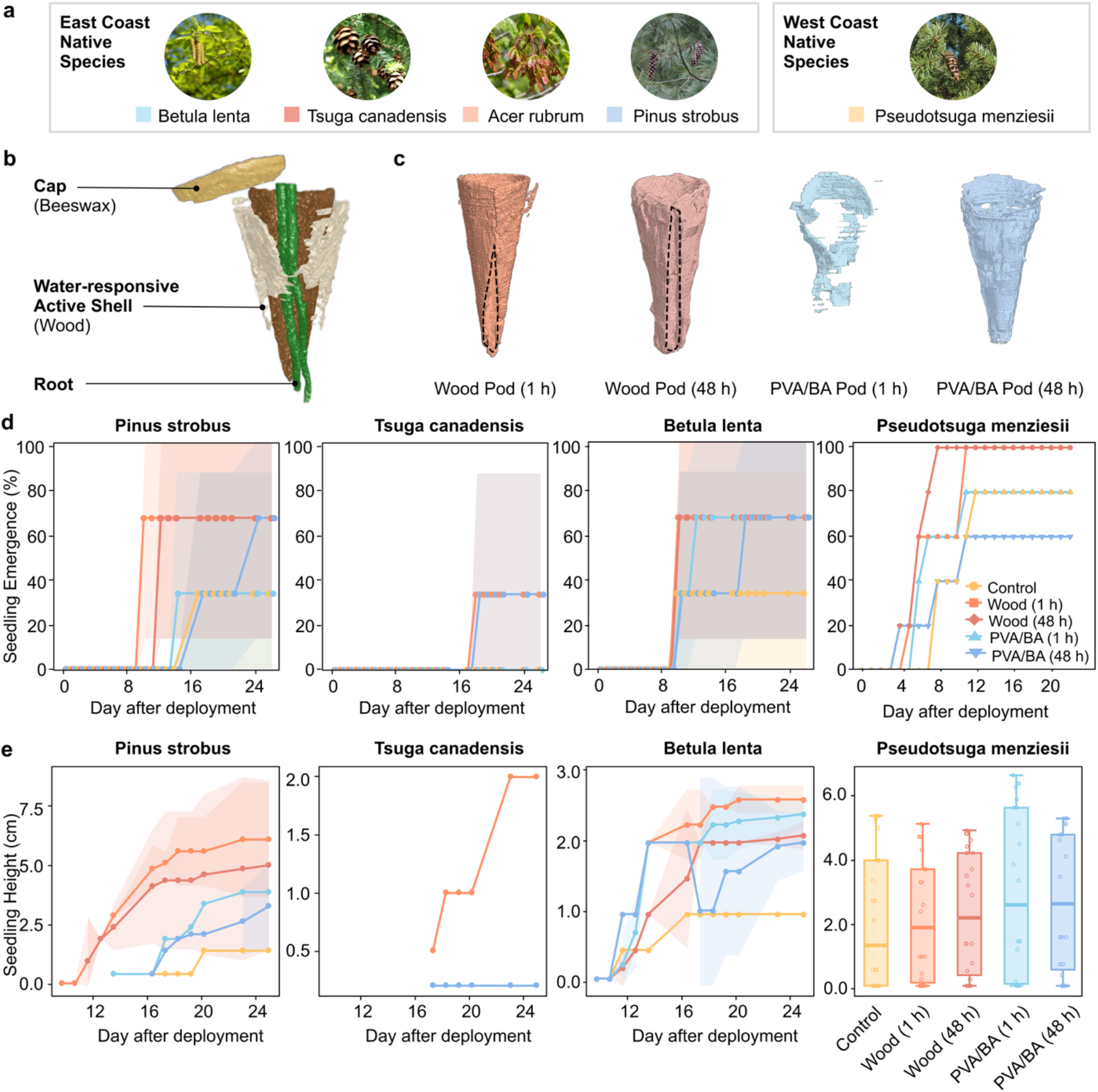
Indoor test for East and West Coast seeds. **a.** East Coast native species (*Betula lenta, Tsuga canadensis, Acer rubrum, Pinus strobus*) and a West Coast native species (*Pseudotsuga menziesii*) used in this study. Color codes are consistent across panels b–g and denote the same seedpod designs: yellow, control (no seedpod); orange, wooden seedpod (1 h); red, wooden seedpod (48 h); sky blue, PVA/BA seedpod (1 h); and blue, PVA/BA seedpod (48 h), where values in parentheses indicate activation delay. **b.** Multilayer wooden seedpod architecture showing the beeswax cap, wax-based delay timer, water-responsive wooden shell, and emerging root. **c.** Representative segmented reconstructions illustrating geometric variability across seedpod configurations. Germination dynamics by species and seedpod design, showing cumulative germination as a function of time since deployment (*Acer rubrum* showed no germination). Error bars and shaded/lightly colored areas show mean ± 95% confidence interval (n = 8 seeds per treatment, per species). **d.** Post-germination height growth over time for germinated seedlings. **e.** Cumulative seedling emergence over time for seeds deployed using wooden and PVA/BA seedpods with different activation delays in comparison with the control. (mean ± s.d., n = 5 per treatment).

The multilayer wooden seedpod is comprised of a beeswax cap and a water-responsive wooden shell that opened to permit root emergence (Fig. 4b). Micro-computed tomography of germinated samples further showed roots extending through the opened pod structures and into the surrounding soil across the different seedpod configurations (Fig. 4c, Supplementary Note 6,7 and Supplementary Fig. 12).

Emergence responses varied among species and seedpod designs (Fig. 4d). The clearest response occurred in *B. lenta* and *P. strobus*, for which wooden seedpods accelerated early emergence, particularly when programmed to open after 1 h (Supplementary Note 8). By week 3, predicted emergence reached ∼62–64% for wooden seedpods (1 h) and ∼60% for wooden seedpods (48 h), compared with ∼26–27% for the direct-sown control. In *P. strobus*, wooden seedpods (1 h) exceeded the control as early as week 1 (Holm-adjusted *p* = 0.025; p-values adjusted for multiple pairwise comparisons), although most other pairwise differences were uncertain after multiplicity correction (Supplementary Fig. 13). A similar pattern was observed in *P. menziesii*: the wooden seedpod (48 h) reached 86% emergence in week 1, compared with 19% in the direct-sown control.

This earlier emergence did not alter subsequent seedling height growth. Over the ∼27-day observation period, neither seedpod design nor its interaction with time significantly affected seedling height. The design p-value indicates the significance of seedpod design effects, while the interaction p-value tests whether the effect of design varies with time (*Betula*: design p = 0.207, interaction p = 0.669; *Pinus*: design p = 0.878, interaction p = 0.303; Fig. 4e). Together, these results show that wooden seedpods can accelerate early emergence in responsive species while maintaining comparable post-germination growth dynamics. Similarly, *P. menziesii* seedling height remained comparable across designs (*p* = 0.367)

Collectively, these experiments demonstrate that programmable activation can accelerate early establishment across morphologically distinct seed types while maintaining baseline post-emergence growth dynamics. The water-responsive wooden shell appears to facilitate timely hydration at the seed-soil interface without imposing detectable growth penalties.

### Field Studies in Northern California Forest

A field test was conducted at the University of California, Berkeley’s Blodgett Forest from May to July 2025. Supplemental irrigation achieved an average of 43% soil moisture (Supplementary Note 9). Coastal Douglas-fir was selected for its importance as a dominant forest component across its native range, including the northern California area where we conducted the field tests. Two spatially separated blocks are used to minimize site-specific variability and reduce potential confounding effects from local environmental conditions. Each block contains two plots for a total of four test sites (Fig. 5a). At each site, five different treatments are deployed: wooden seedpods (1 h and 48 h), PVA/BA seedpods (1 h and 48 h) and control (seed-only). Each seedpod contained three *P. menziesii* var. *menziesii* seeds. A total of 150 seeds were deployed across all sites. Control groups consist of 150 bare seeds placed directly into 5 cm-deep depressions (Fig. 5b).

**Figure 5.**
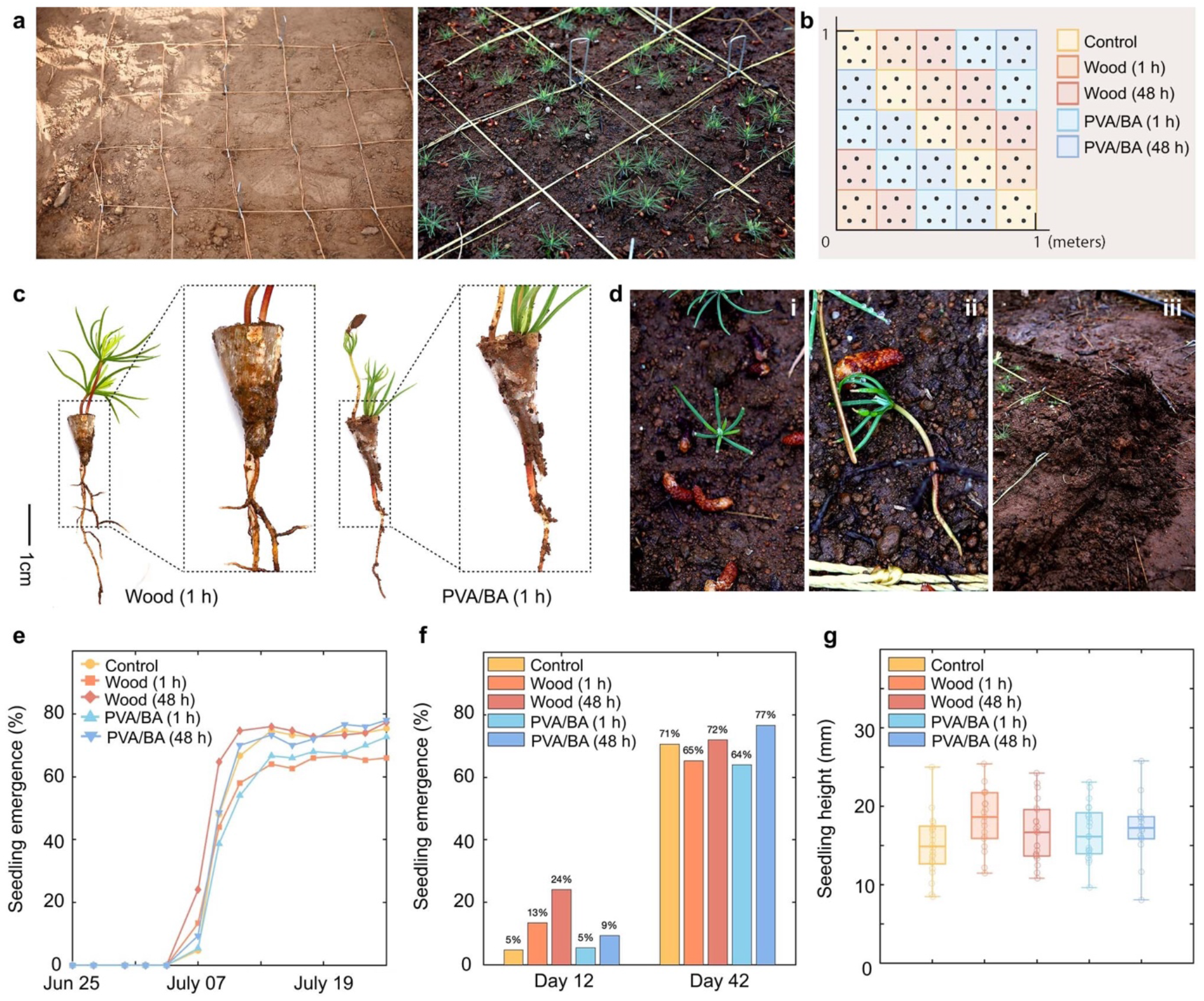
Outdoor testing for the field studies. **a.** Field study plot with artificial irrigation and fencing system on the first day and final day of observation, showing emerged seedlings. **b.** Randomized plot design for each treatment to minimize location-specific effects. **c.** Root extraction and analysis across treatments: the wooden seedpods (1 h) and the PVA/BA seedpods (1 h). **d.** Impacts of the wildlife: (i) foliar herbivory, (ii) uprooting, (iii) tunneling damage at plot edges. **e.** Germination rates over the full observation period (∼4 weeks). Each seedpod contained 3 seeds (n = 50 seedpods, 150 seeds per treatment). **f.** Seedling emergence (%) from the first day of emergence to the final observation. **g.** Stem length for each treatment after the extraction (mean ± s.d., sample sizes: control, n = 22; wooden seedpods (1 h), n = 19; wooden seedpods (48 h), n = 23; PVA/BA seedpods (1 h), n = 22; PVA/BA seedpods (48 h), n = 16.

Germination was monitored at 3-day intervals over a one-month period. Post-extraction analysis showed that root-to-stem ratios are consistent across treatments, indicating that seedpods do not hinder germination (Fig. 5c). Wildlife disturbances that reduce germination rates beginning at day 20 included foliar herbivory (Fig. 5d-i), uprooting (Fig. 5d-ii), and tunneling at plot edges that disrupted soil structure (Fig. 5d-iii). However, no discernible negative impacts of any treatment on overall seedling emergence or growth were observed when compared to the control. Several treatments improved early emergence compared to the control, with the wooden seedpod (48 h) achieving the highest early emergence rates relative to all other treatments (Fig. 5e), consistent with the indoor test results.

All treatments except PVA/BA seedpod (1 h) promoted faster emergence than that of the control (Fig. 5f). The first observation of seedling emergence occurred on Day 12 following seed deployment, at which time the greatest proportion of deployed seeds that emerged as seedlings (24%) was observed in the wooden seedpod (48 h). This treatment maintained an emergence advantage through the final recording on Day 42. Compared to control, both wooden and PVA/BA seedpods showed comparable or higher final emergence (64-77%). Rapid germination and emergence are desirable because they (1) allow practitioners to align seed deployment with weather conditions favorable for germination, and (2) help seedlings transition quickly beyond the vulnerable seed stage, reducing granivore risk.

## Discussion

Ecological restoration is constrained not only by getting seeds buried into the ground but also by what happens to them after dispersal. Here we show that biodegradable seedpods regulate the post-dispersal environment through both spatial (microclimate) and temporal (activation timing) controls, altering germination timing and allowing for early establishment without restricting root emergence or penalizing subsequent growth. Rather than acting as passive delivery devices, the seedpods function as active regulators of seed–soil interactions, demonstrating that the post-dispersal environment can be intentionally engineered. Compared to other seed enhancement coating techniques, the pods introduced here can gate the microclimate around the seeds for a preprogrammed duration.

A second contribution is methodological. Because the pod encloses an air volume, sensors can be directly embedded inside so that the seeds’ microenvironment can be directly measured rather than inferred from downstream plant performances like how other seed enhancement technologies were typically evaluated. This turns the post-dispersal microclimate from an assumed mechanism into a measured variable and makes it possible to investigate which conditions matter and for how long, rather than only whether a treatment worked.

What our pod does to the seed is the warmer, drier cavity conditions and the timed opening, which likely account for the observed differences in emergence and seedling performance. Elevated temperature would advance the thermal accumulation required for germination, while the delayed opening postpones full soil contact until wetting is sustained. The contrasting seedpod opening strategies, crack-open wooden and dissolution-open PVA/BA, hold unique distinctions in protection, exposure timing and residue. Rather than aiming for a universally optimal design, we treat the contrast as a design space and suggest that different post-dispersal strategies may be suited to different restoration objectives, paralleling the diverse environmental regulation achieved by natural seed structures.

Several limitations bound the conclusions. We tested a limited number of species at a single field site. And because the field plot was irrigated, the gating behavior has not yet been tested against the natural rainfall variability it was designed for. Our outdoor monitoring effort was stopped after a month, so the longer-term seedling growth is unknown. Manufacturing throughput and per-unit cost remain far from what landscape-scale deployment requires. In addition, the pods used for microenvironment monitoring were larger than those used in germination tests, in order to accommodate the temperature and humidity sensors, and the effect of pod size was not studied systematically.

Within those bounds, the results address a long-standing challenge in forest restoration. Direct seeding has low yields even at high sowing rates because even we can let the seeds reach a site and ideal soil depth, we cannot ensure a suitable establishment condition. Warmer and more variable precipitation is narrowing the window in which a first-year seedling can survive to its first summer, as a seed that germinates during a false-start rain is simply lost later. Our approach suggests that suitable environment and microsite need not be naturally found, some of its properties such as suitable temperatures and moisture conditions can be manufactured, and its timing to germinate can be set by the practitioner rather than by the seed’s own dormancy threshold.

This extends restoration technology from delivering seeds to regulating the conditions they encounter. Through treating a seed’s first weeks in the soil as a designed and measured condition, we hope to give restoration ecology an experimental handle on a stage of establishment it has largely had to infer.

## Methods

### Materials

Polyvinyl alcohol (PVA; molecular weight 13,000-23,000 g/mol; 87-89% hydrolyzed) was purchased from Sigma-Aldrich. Boric acid (BA; DNAse-, RNAse-, and protease-free, 99.5%) was purchased from Thermo Scientific. All chemicals were used without further treatment. For the PDMS mold, Sylgard 184 silicone elastomer (Dow Inc., USA), consisting of a dimethylsiloxane-based prepolymer and a curing agent, was used. Flat-sawn maple wood veneers (VeneerSupplies.com), with a thickness of 0.4 ± 0.05 mm were utilized. Beeswax pellets (triple-filtered, cosmetic grade; Beesworks) and carnauba wax flakes (natural grade; TooGet) were purchased from Amazon (USA).

### Chemical wash

Wood veneer strips (0.45 mm thickness, 2.2 g) were boiled for 5 minutes in a 100 g solution consisting of 10 wt% sodium hydroxide (Bell Chemical). The strips were removed from the solution and rinsed with hot water and air-dried completely before use.

### Water shock

Prior to forming, the strips were rehydrated by submerging in water for 3 min, followed by wiping off excess surface moisture and allowing them to partially dry for approximately 2 min.

### Preparation of PVA/BA solutions

PVA was dissolved in deionized (DI) water to form a homogeneous solution. With stirring, BA solids were added directly, resulting in the immediate formation of white precipitates. Continuous stirring at 80 ℃ redissolved the precipitates, yielding a homogeneous solution. The total weight fractions of PVA and BA in water were 10, 15, and 25 wt%, respectively. For example, to prepare a PVA/BA (30:1) solution, PVA (4.84 g) was dissolved in DI water (45 g). BA (0.161 g) was then added with vigorous stirring, forming instant white precipitates. The mixture was stirred for an additional 2 h at 80 ℃ until the precipitates were completely dissolved, producing a homogeneous solution.

### Fabrication of PDMS molds

Sylgard 184 silicone elastomer base and curing agent were mixed in a 10:1 weight ratio and cast onto a 3D-printed polylactic acid (PLA) master. The master, fabricated using a Bambu Lab X1C 3D printer, consisted of a 3 × 5 array of cones, with each cone having a diameter of 8.4 mm and a length of 20 mm. After curing at room temperature for 48 h, the PDMS mold was carefully peeled off.

### Fabrication of the PVA/BA seedpods

The PVA/BA solution was injected into the PDMS mold using a transfer pipette and left to evaporate for 7 days to form a conical shell with a high-fidelity sharp tip. The dried seedpods were then carefully removed from the mold using a pair of tweezers.

### Fabrication of the wax coating solution

Beeswax pellets and carnauba wax flakes were mixed in IPA and heated under magnetic stirring using a double-boiler setup in a covered vessel. The temperature was gradually increased to 90 °C to obtain a homogeneous solution and then reduced to 60 °C prior to coating.

### Application of the wax coating

The PVA/BA seedpods were immersed in the wax-IPA solution at 60 ℃ for 3 s, ensuring only the outer surface was in contact with the solution, followed by air-drying at room temperature for 3 days. 60 ℃ was chosen to be above the melting point of beeswax and below that of carnauba wax. Maintaining this constant temperature throughout all experiments ensured a homogeneous molten blend with stable viscosity, thereby enabling reproducible coating thickness.

### Activation time experiments

Seedpod activation was tested in soil maintained at a gravimetric water content of 43–46% (Supplementary Fig. 8 and Supplementary Note 3). Experiments were conducted at an ambient temperature of 22 °C and relative humidity of approximately 50%. Activation time was defined as the time required for the wooden seedpod to develop a 1 mm opening along the seam or for the PVA/BA seedpod to disintegrate. Wooden and PVA/BA seedpods were tested with different wax concentrations, numbers of outer coating layers and inner-coating configurations.

### Diffusion modelling

Activation timing is analyzed using a one-dimensional diffusion model in which water transport through the wax coating sets the activation timescale (Supplementary Note 4). An effective wax diffusivity *D*_w*ax*_ is extracted from gravimetric sorption on coated specimens (Fig. 3d). For wooden seedpods, the wood absorbs water far faster than the wax delivers it, so the activation time is predicted from the wax permeation lag,

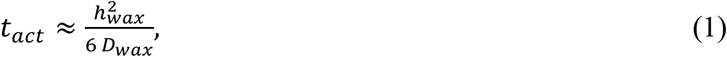

where ℎ_w*ax*_ is the wax coating thickness. For PVA/BA seedpods, disintegration is driven by accumulated hydration damage. We model this with a wax/PVA bilayer diffusion equation coupled to a damage variable above a hydration threshold (Supplementary Note 4, Eq. 4), with parameters fit to the disintegration times in Fig. 3g.

### Soil preparation for performance evaluation

Approximately 85 g of natural forest soil, sourced from the University of California, Berkeley’s Blodgett Forest, CA, was placed into a 180 g specimen cup equipped with five 3/16” diameter drainage holes. To simulate post-precipitation soil moisture conditions, approximately 46.6 g of water was added and allowed to drain, resulting in a gravimetric water content of 43-46%.

### Soil evaluation for indoor tests

For the indoor seedpod tests, a well-drained fertile Arbordale fine sandy loam was used (pH 7.1, organic matter 3.6%) (see Supplementary Note 5).

### Indoor environmental conditions

Northeastern species seed deployment and incubation were conducted in the Guterman Bioclimatic Laboratory greenhouse (Cornell University), where daytime temperature was 73-77 °F (22.8-25.0 °C) and nighttime temperature was ∼70-75 °F (21.1-23.9 °C). Natural ambient lighting was supplemented with advanced LED lighting to provide full-spectrum lighting conditions.

### Indoor test seedling height analysis

Post-germination height trajectories were analyzed using generalized least squares models with first-order autoregressive (AR(1)) errors to account for within-trajectory temporal correlation. Analyses were restricted to successfully germinated seedlings, excluding pre-emergence zeros. For the Northeastern experiment, species with negligible germination were excluded from height modeling.

Fixed effects included centered time (weeks since germination), seedpod design, and their interaction. Significance of fixed effects was evaluated using Type III tests^30–33^. Treatment differences were summarized using estimated marginal means for week-specific height comparisons and growth-rate (slope) contrasts versus the Control with multiplicity adjustment^34^.

### Seed preparation for indoor and field trial

Before deployment of seeds in our soil and seed carriers, we conducted an extensive review of the literature and complementary experimentation to determine appropriate stratification and germination treatments for each tree species. Reliable information on germination rates and handling protocols for temperate tree species is often scattered or incomplete, requiring us to compile data across multiple sources and adapt it through preliminary trials^35–38^. We standardized our approach to a 24-hour soak in a solution of lukewarm water and gibberellic acid (GA₃ ; 0.167 mg/mL, prepared as 20 mg in 120 mL), followed by cold stratification (3–5 °C) for approximately 30 to 60 days, depending on the species. This preparatory work ensured that seeds were synchronized for timely germination and allowed us to establish consistent starting conditions across species. For the control treatment (seed-only), seeds were planted at a depth approximately twice the seed diameter.

### Field trial site preparation and irrigation system

The field trial was conducted on the grounds of the University of California, Berkeley’s Blodgett Experimental Forest (38.907815 N, -120.673146 W; approximately 1200 m elevation), within 100 m of the main research station building. The forest is mixed-conifer with major components of Ponderosa Pine, Sugar Pine, Incense Cedar, White Fir, and Douglas-fir. Two plots with an area of approximately 3 m^2^ were established one meter apart at the base of a mature Douglas-fir, and duff and litter were scraped away to expose mineral soil in each plot. Plots received direct sunlight for most of the day. An automated overhead spray irrigation system was designed to deliver sufficient soil moisture for seed germination, seedling emergence, and survival through the trial period. Irrigation was achieved with four Tempo© FSA24-Q Jet Stake assembly 90-degree spray emitters established along the perimeter of each caging exclosure. Emitters were operated with a Hunter© BTT-100 flow controller programmed to irrigate in cycles of five minutes on, 20 minutes off, from 8:30 a.m. to 5 p.m. daily. Water was applied at 1.64 L/m^2^ per five-minute watering cycle, with a total of 32.8 L/m^2^ water applied daily to each plot.

### Micro-computed tomography

We studied the seedpod and seed interaction with the soil and each other, by acquiring X-ray micro-computed tomography (μ-CT) scan images with a SkyScan1276 micro-CT scanner at 20 to 40 µm voxel resolution. The CT-scans were performed at four different time points. These scan intervals were t = 0 h with no additional watering, t = 1 h with the first watering applied, t = 24 h with maximum water holding capacity (MWHC) maintained, and t = 1 week at MWHC. We captured the seedpod decomposition, seedling emergence, and growth by manually segmenting and measuring the temporal changes in seedpod opening metrics in Object Research Systems (ORS) Dragonfly 3D World (Comet Technologies Canada Inc. (2025)).

## Data availability

Data generated and analyzed during the study will be available at Zenodo. Correspondence and requests for materials, digital models, processing protocols and datasets should be addressed to.

## Supporting information

S1_Moisture-induced crack-open seedpod

S2_Moisture-induced opening of Erodium seedpod

## Acknowledgements

We thank Ariel Thomson Roughton, Amy Mason and Rob York for supporting the field test at Blodgett Forest Research Station and Ramona Werner for insightful suggestions. The authors acknowledge funding support from the U.S. National Science Foundation, including NSF-GCR-2428641 (L.Y., T.Z., S.Y., T.B.), IIS-CAREER-1847149 (L.Y.), and OIA-2428643 (T.Z.).

## Author contributions

L.Y., S.Y., S.Y., and Y.H. conceived the initial concept. L.Y., T.Z., S.Y., T.B. and K.S. supervised the project. L.Y., S.Y., Y.H., and F.J. wrote the manuscript. S.Y. performed mechanical and material characterization and microenvironment monitoring tests. S.Y. and Y.H. performed fabrication and performance evaluation. Z.Z. supported the development of wax coating methods on PVA/BA. T.B. and F.J. performed the indoor test and micro-CT scanning and conducted statistical analysis of indoor test results. L.Y., S.Y., M.F., and E.M. conducted field test. Y.Y. performed soil characterization for the Berkeley Forest Blodgett Forest. H.L. reviewed the manuscript draft and provided feedback. S.Y., T.Z., T.B., K.S. and L.Y. provided scientific and experimental advice. All authors commented on the manuscript.

## Competing interests

There are no competing interests.

## Notes

### Competing Interest Statement

The authors have declared no competing interest.

