## Supplementary Material for "Hygrochastic Seedpods: Engineering Microclimates for Forest Restoration"

### Supplementary Information

**This PDF file includes:**

Supplementary Notes 1 to 9

Supplementary Figures 1 to 13

#### Supplementary Note 1. Environmental Triggers of Seed Release.

Evidence from tropical to temperate ecosystems indicates that many species time seed release to coincide with periods of moisture availability, facilitating rapid seed germination and desiccation avoidance[11–15] . For example, a field study of 82 species native to seasonally dry Brazilian savanna found that species with seeds that were released from fruit capsules at the onset of the rainy season exhibited less dormancy than species that released seeds at other times of the year[8]. In a field study of environmental triggers of seed release in black cottonwood (*Populus balsamifera* ssp. *trichocarpa*) trees in southeastern British Columbia, Canada, Herbison et al. (2015) found that 18 seed release events spanning four years all immediately followed rain events and concluded that rain is the primary trigger for seed release in black cottonwood. Rain events that led to seed release from catkins provided soil moisture conducive to seedling establishment, whereas cottonwood seedling survival can be strongly limited by dry soil conditions[13].

#### Supplementary Note 2. Material selection and fabrication of seedpods

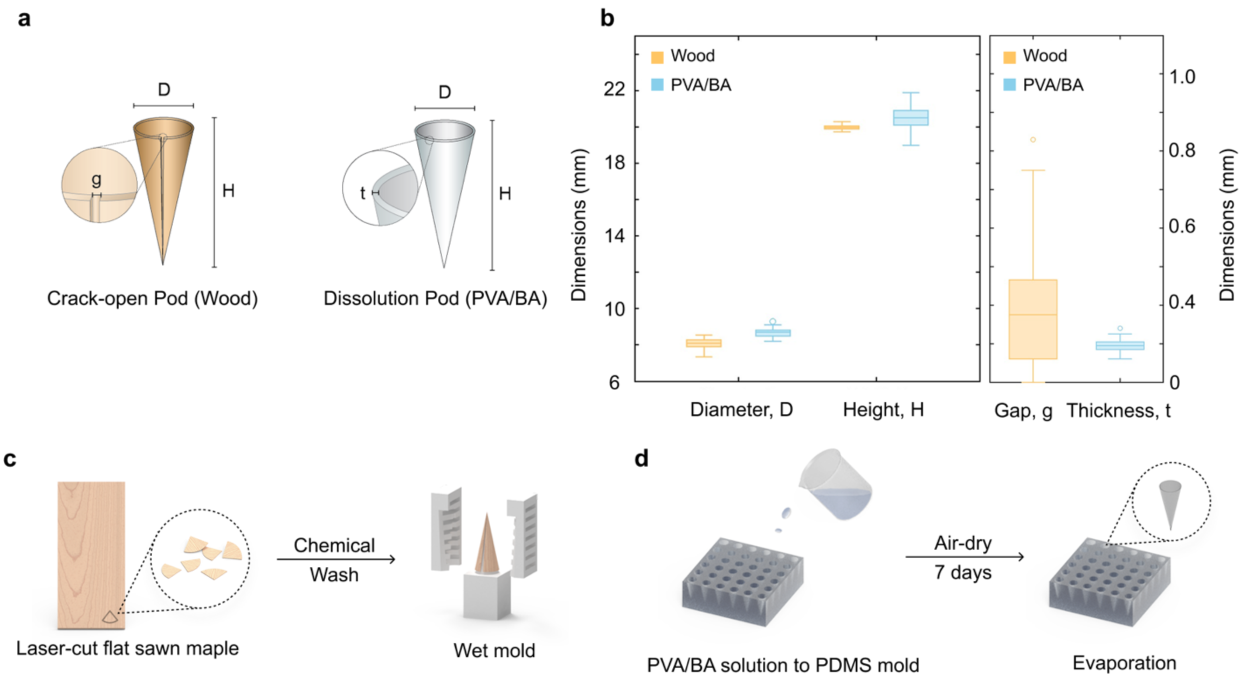


**Supplementary Figure 1 | a.** An illustration of a wooden and a poly(vinyl alcohol)/boric acid (PVA/BA) seedpod with key measurements. **b-c.** Fabrication process of wooden seedpods **(b)** and PVA/BA seedpods **(c)**, **d.** with measured dimensions from 100 fabricated seedpod samples.

To develop a seedpod that dehisces in humid conditions, the selected material must be hygromorphic and capable of forming a cone shape for effective deployment. For the crack-opening seedpods, natural maple wood is used owing to its humidity-driven deformation. Its diffuse-porous anatomy provides uniform mechanical properties (elastic modulus, E~12.6 GPa)[39] along its grain direction, and easy manufacturability. The 0.45 mm-thick veneer sheets are laser-cut with the grain oriented parallel to the cone axis to facilitate molding, and then chemically washed with 10wt% sodium hydroxide solution (see Methods and Supplementary Fig. 1c).

For the water-soluble seedpods, PVA is selected, which can be crosslinked by BA forming boronate esters upon drying to form a solid, rigid film by simply casting the mixed aqueous solution[22], or molded into cones with excellent reproducibility and scalability for mass production (Supplementary Figure 1). Upon exposure to water (pH ≤7), boronate esters (pKa of

BA, 9.2) can easily dissociate, leaving PVA/BA dissolved. To optimize the interplay between soil interaction, dissolution rate, seedpod stiffness, and strength, we varied the molecular weights of PVA (13,000-23,000 g/mol) and PVA/BA mass ratios with a PVA/BA ratio of 1:30 (wt/wt) to achieve optimal performance (see Methods and Supplementary Fig. 1d). This crosslinking improves structural stability while preserving the environmental responsiveness, ensuring that the seedpods retain their shape during storage, handling, and deployment yet undergo programmed dissolution upon exposure to soil moisture.

For the tailored manufacturing process, first, we laser-cut 0.45 mm thick wood veneer into flattened cone shapes and chemically wash with a 10 wt% sodium hydroxide aqueous solution to increase the compliances of wood for moldability (Method, Supplementary Fig. 1). The delignified veneer is then rehydrated, mechanically clamped between a perforated inner-outer mold, and pulled into a tight conical shell, yielding a reproducible shell geometry with a tight curvature without cracking (Fig. 2c). In parallel, the PVA/BA aqueous solution is cast into a polydimethylsiloxane (PDMS) cone mold (Fig. 2d, Supplementary Figs. 3 and 4). Slow solvent evaporation produces thin, uniform PVA/BA shells conforming to the same conical geometry as wooden seedpods. The samples are mass-produced with key dimensions monitored as quality-control metrics to assess fabrication repeatability (Fig. 2e, n = 100 for each seedpod type). Wood and PVA/BA seedpods each offer complementary advantages. Wooden seedpods can be formed relatively quickly from natural veneers, whereas PVA/BA seedpods require a slower drying process but achieve much higher geometric consistency.

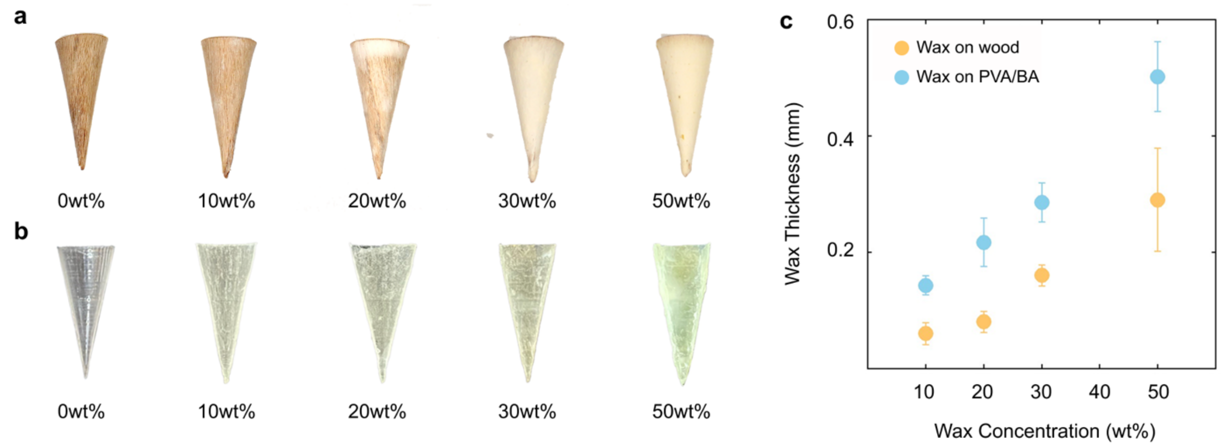

**Supplementary Figure 2 | a-b.** Photos of wood (**a**) and PVA/BA (**b**) seedpods coated with wax of various concentrations. **c.** Wax coating thickness as a function of the concentrations of carnauba wax and beeswax (1:1 wt/wt) in IPA.

To precisely control the opening time of the seedpods, we dip-coat the seedpod with a thin layer of wax (Supplementary Fig. 2a,b) blended from carnauba wax and beeswax in equal mass to balance their mechanical and thermal properties (Supplementary Fig. 6). Carnauba wax has a higher flexural modulus (~536.5 ± 56.1 MPa) and melting temperature ($T_m$ ~85°C), which provides structural protection to the seedpod tips[40–44]. However, its brittleness may cause it to fracture under localized stress. In contrast, beeswax has a lower flexural modulus (~75.1 ± 11.1 MPa) and offers greater viscoelasticity and flexibility, but its relatively low $T_m$ (~65°C) makes it prone to be softened or deformed at elevated storage or field temperatures [42,45,46]. By combining these two waxes, we create a composite coating that balances stiffness (~268.1 ± 16.4 MPa), toughness, and hydrophobicity, thereby serving as an effective moisture diffusion barrier (Supplementary Fig. 7). To ensure uniform wetting and controlled film formation, we adopt isopropyl alcohol (IPA) as a solvent for wax to reduce surface tension. We vary the concentration of the wax solutions to tune the coating thickness, thereby further regulating the hygroscopic response (Supplementary Fig. 2c). The thicker wax coating increases resistance to water diffusion, delaying seedpod opening time.

**Supplementary Note 3. Microenvironment Characterization of Seedpods.**

To understand the microenvironment within the soil and inside two types of seedpods (wooden and PVA/BA seedpods), we conducted a 48-hour experiment where we let the soil air-dry. The soil was first saturated to full water-holding capacity using a self-watering system[23], and then the water source was removed before deploying the seedpods. Temperature and relative humidity were continuously measured both in the surrounding soil and within the interior of the seedpods. Due to the physical size of the BME280 sensor, the experimental sample volume was increased by approximately sevenfold to accommodate sensor placement.

The results show that both seedpod types create distinct microenvironments relative to the surrounding soil. The wooden seedpods exhibited a temperature increase of 2.513 ± 0.748 °C and a relative humidity decrease of −16.910 ± 2.925% (Supplementary Fig. 9), while the PVA/BA seedpods showed a smaller temperature increase of 2.010 ± 1.165 °C and a relative humidity decrease of −10.129 ± 6.026% (Supplementary Fig. 10). These findings suggest that the wooden seedpod provides a drier and warmer internal environment compared to the PVA/BA seedpods under the same drying conditions.

These microenvironmental differences suggest distinct implications for germination. The slightly elevated temperature within both seedpods may promote metabolic activity and accelerate early germination processes at a time when water is sufficiently accessible to enable successful seedling emergence and establishment. At the same time, the reduced relative humidity, particularly in the wooden seedpods, indicates a drier environment compared to the surrounding soil, which may promote favorable water potential levels without excess moisture. As a result, both seedpods may create conditions that support modest seed warming without undue impact on seed hydration, potentially enhancing metabolic activity leading to early emergence as compared to seeds in bare soil.

**Supplementary Note 4. Activation time prediction.**

Activation time is the time required for a wooden seedpod to open to a 1 mm gap through the seam, or for a poly(vinyl alcohol)/boric acid (PVA/BA) seedpod to disintegrate. Both seedpod types share the same transport geometry, in which a wax coating covers an active substrate (wood or PVA/BA) that encloses a closed seed cavity. The reference problem is one-dimensional Fickian diffusion through this wax/substrate bilayer, with $C(0,t) = C_s$ at the immersed outer surface, flux continuity at the wax/substrate interface, and $\partial C/\partial x = 0$ at the inner substrate surface. The inner condition treats the closed cavity as a no-flux boundary, neglecting small contributions from water vapour transfer to cavity air and absorption by the seeds, which are negligible compared with the wax-side influx over the activation timescale. The wood and PVA/BA analyses below are simplified reductions of this reference problem, chosen to match the dominant physics of each case.

The effective wax diffusivity $D_{wax}$ was measured by gravimetric sorption on wax-coated wood-shell specimens of thickness $h$. Specimens were fully dried and weighed to obtain the initial mass $M_0$. Specimens were then immersed in water, periodically removed, gently blotted to remove surface water, and reweighed to obtain the time-dependent mass $M_t$ until the mass reached a plateau $M_\infty$. Normalized uptake $C_t = (M_t - M_0)/(M_\infty - M_0)$ varied linearly with $\sqrt{t}$ at short times (mean ± s.d., $n = 5$, Fig. 3g). The slope $\theta$ of the initial linear region was used to compute $D_{wax}$ via the standard short-time Fickian approximation[47]:

$$D_{wax} = \pi \left(\frac{h\,\theta}{4}\right)^2. \tag{2}$$

For the wooden seedpod, water diffusivity in wood is several orders of magnitude larger than in the wax coating ($D_{wood} \sim 10^{-10}$ m² s⁻¹ versus fitted $D_{wax} \sim 10^{-13}$ m² s⁻¹)[48]. The wood absorbs water much faster than the wax delivers it, so the interfacial concentration on the wood side stays well below saturation throughout the activation period. The wax/wood interface is therefore treated as a perfect sink ($C \to 0$ on the wax side), reducing the bilayer to the classical Daynes–Barrer permeation problem for the wax layer alone. Activation time is then approximated by the wax permeation lag,

$$t_{act} \approx \frac{h_{wax}^2}{6\,D_{wax}}. \tag{3}$$

The time required to fill the wood, $\sim h_{wood}^2/D_{wood}$, is on the order of minutes for the 0.45 mm wood shell used here, compared with wax lag times of tens of hours, and is omitted. Equation (3) is consistent with the quadratic dependence of $t_{act}$ on coating thickness in Fig. 3d, e.

For the PVA/BA seedpod, disintegration occurs by accumulated hydration damage rather than by a single fracture event. The activation criterion is therefore a time integral of the local concentration field rather than a threshold concentration, and the bilayer is solved numerically with $D_{wax}$ in the wax layer, an effective $D_{PVA}$ in the hydrated PVA/BA, and the boundary conditions stated above. A damage variable $d \in [0,1]$ is integrated on top of the resulting concentration field at a rate

$$\frac{\partial d}{\partial t} = k_{dmg}\max\left(\frac{C - C_{crit}}{1 - C_{crit}}, 0\right)(1-d), \tag{4}$$

where $C_{crit}$ is the hydration threshold below which no damage accumulates. Disintegration is the time at which the spatial average of $d$ first reaches $d^* = 0.80$. The diffusion and damage are coupled in one direction only. $C(x,t)$ is obtained from the diffusion equation and drives $d$ through equation (4), but $d$ does not feed back on $D_{PVA}$, which is held constant. Hydration would soften the polymer network and increase its permeability. Here the constant-$D_{PVA}$ assumption absorbs this into an effective diffusivity averaged over the disintegration interval. Equations were integrated using a one-dimensional finite-volume scheme with backward Euler time stepping.

Four parameters were fit simultaneously to the six conditions in Fig. 3f: two values of $D_{wax}$ (one per wax concentration, shared across PVA/BA compositions) and two values of $k_{dmg}$ (one per PVA/BA composition). Wax thicknesses were measured directly ($h_{wax} = 0.286 \pm 0.038$ mm at 30 wt% and $0.502 \pm 0.067$ mm at 50 wt%, $n = 5$). PVA/BA shell thicknesses were 200 μm at 10 wt% and 250 μm at 15 wt%. The hydration threshold $C_{crit}$ was fixed at 0.45. The two wax-free conditions anchor $k_{dmg}$ for each PVA/BA composition independently of the wax-coated conditions, so the wax-coated fit effectively tests two free parameters against four measurements. Fitted parameter values are reported in Supplementary Table S4.1, and per-condition predictions in Supplementary Table S4.2.

**Supplementary Table S4.1.** Fitted parameters at $d^* = 0.80$.

| Parameter | Value |
|---|---|
| $D_{wax}$ (30 wt% wax) | $3.40 \times 10^{-13}$ m² s⁻¹ |
| $D_{wax}$ (50 wt% wax) | $4.98 \times 10^{-13}$ m² s⁻¹ |
| $k_{dmg}$ (10 wt% PVA/BA) | $1.71 \times 10^{-4}$ s⁻¹ |
| $k_{dmg}$ (15 wt% PVA/BA) | $1.66 \times 10^{-4}$ s⁻¹ |

**Supplementary Table S4.2.** Measured and predicted disintegration times for the six conditions in Fig. 3g.

| PVA/BA (wt%) | Wax (wt%) | $h_{wax}$ (µm) | Measured (h) | Predicted (h) | Residual |
|---|---|---|---|---|---|
| 10 | 0 | — | 3.0 | 2.9 | −4% |
| 10 | 30 | 286 | 61.8 | 72.2 | +17% |
| 10 | 50 | 502 | 102.1 | 108.5 | +6% |
| 15 | 0 | — | 3.0 | 3.1 | +4% |
| 15 | 30 | 286 | 110.8 | 81.2 | −27% |
| 15 | 50 | 502 | 127.4 | 118.3 | −7% |

Predictions agree with measurements to within ~30% across all six conditions, with the largest residual (−27%) on the 15 wt% PVA / 30 wt% wax point. Fitted $D_{wax}$ values lie within the literature range for water diffusivity in wax films[39,41,43]. The two values differ by a factor of 1.5 between coatings, while $h_{wax}^2$ differs by a factor of 3.1, so coating thickness controls disintegration timing more strongly than the variation in fitted diffusivity. The 1.5× change in fitted $D_{wax}$ likely reflects microstructural differences (porosity, crystallinity, morphology) between the two coatings rather than a change in molecular-scale water diffusivity in wax.

The choice $d^* = 0.80$ defines disintegration but is not derived from a measurement. Repeating the fit with $d^*$ varied from 0.50 to 0.95 (Supplementary Table S4.3) leaves the fitted $D_{wax}$ values unchanged to three significant figures and the per-condition residuals unchanged, while $k_{dmg}$ rescales to compensate for the threshold shift. The wax permeation parameters are therefore robust to the choice of $d^*$. The absolute value of $k_{dmg}$ is not, and should be interpreted in conjunction with the threshold.

**Supplementary Table S4.3.** Fitted parameters as a function of damage activation threshold.

| $d^*$ | $D_{wax}$ (30 wt%) | $D_{wax}$ (50 wt%) | $k_{dmg}$ (10 wt%) | $k_{dmg}$ (15 wt%) |
|---|---|---|---|---|
| 0.50 | $3.39 \times 10^{-13}$ | $4.97 \times 10^{-13}$ | $7.4 \times 10^{-5}$ | $7.2 \times 10^{-5}$ |
| 0.65 | $3.39 \times 10^{-13}$ | $4.98 \times 10^{-13}$ | $1.12 \times 10^{-4}$ | $1.09 \times 10^{-4}$ |
| 0.80 | $3.40 \times 10^{-13}$ | $4.98 \times 10^{-13}$ | $1.71 \times 10^{-4}$ | $1.66 \times 10^{-4}$ |
| 0.90 | $3.40 \times 10^{-13}$ | $4.99 \times 10^{-13}$ | $2.42 \times 10^{-4}$ | $2.36 \times 10^{-4}$ |
| 0.95 | $3.41 \times 10^{-13}$ | $4.99 \times 10^{-13}$ | $3.11 \times 10^{-4}$ | $3.05 \times 10^{-4}$ |

Our models have a few simplifications. The flat-slab geometry neglects cone curvature, taper, and seam effects. Equation (3) assumes wood water-uptake is fast on the wax permeation timescale, which holds for the coatings tested here but would break down for sufficiently thin wax layers. Equation (2) treats $D_{wax}$ as an intrinsic wax property and applies it to coatings of arbitrary thickness $h_{wax}$ on intact seedpods. Equation (4) holds $D_{PVA}$ constant during integration and so neglects feedback from accumulated damage on polymer permeability; a more mechanistic treatment would couple $D_{PVA}$ to $d$ and solve the resulting nonlinear system. The no-flux inner boundary condition is an approximation as noted above. These simplifications affect quantitative accuracy but not the wax-thickness control of activation identified here.

**Supplementary Note 5. Soil analysis results from Cornell orchards.**

The soil type is an Arbordale fine sandy loam (0–3% slopes, ArB), a fertile, moderately well-drained, medium-textured soil with low clay content.

| Parameter | Value | Interpretation |
|---|---|---|
| Soil pH | 7.1 | Normal (OK) |
| Phosphorus (P) | 18 lbs/acre | High - No added P needed |
| Potassium (K) | 305 lbs/acre | High - No additions needed |
| Organic matter | 3.6% | OK |
| Nitrogen (N) | Only nitrogen might be needed | Apply if needed |

**Supplementary Note 6. Study of seedpod-soil-environment interaction at indoor tests.**

Prior to deployment of the seedpods, we sieved (4 mm) and homogenized Cornell Orchards soil. For easy handling and X-ray CT imaging the Cornell Orchards soil was filled into 120 mL MED PRIDE Disposable Specimen Cups as containers to hold four seeds of the same tree species in four different seedpod designs and a Control (no seedpod). Water drainage from the specimen cups was facilitated by drilling five 3/16" diameter holes into the bottom of each cup. We deployed four different seedpod designs (Wood (1 h), Wood (48 h), PVA/BA (48 h), PVA/BA (1 h)) after settling the soil by jolting the filled specimen cup on a table. To standardize soil moisture across replicates, each specimen cup was filled with Cornell Orchards silt loam to about the 100 mL mark (initial gravimetric water content ≈ 30%) and watered with 34 mL using a calibrated spray bottle (30 strokes). After infiltration and drainage, cups were weighed following a 2–3 h equilibration period to determine stable wet weight. This established the maximum water holding capacity (MWHC) of the soil between 35 and 41% gravimetric water content. Daily watering was subsequently adjusted based on weight loss to maintain soil moisture close to this target, with a minimum of four spray strokes per cup to ensure even distribution among seedpods. This procedure gave us a good idea about the decomposition of the seedpods under saturated soil moisture conditions. We also deployed *Pseudotsuga menziesii* seeds from our collaborators in California in four different seedpod designs (PVA/BA (1 h); PVA/BA (48 h); Wood (1 h); Wood (48 h)) and achieved a stable gravimetric soil water content ($\theta m$) of 43–46% in Berkeley Forest soil, applying the same procedure as for the Northeastern species.

#### Supplementary Note 7. Indoor test seedling emergence analysis.

Statistical analysis was performed using R for statistical computing (R version 4.5.2, 2025-10-31; R Core Team (2025)), and the significance threshold was set to $\alpha = 0.05$. Daily emergence was coded at the specimen cup × seedpod design level as a cumulative binary outcome (0/1), defined by the first observation of at least one germinated seed per specimen. Analyses were conducted separately for each experiment, with species included as a factor. The Northeastern species experiment included 8 replicates (specimen cups), and the Douglas fir experiment included 5 replicates (specimen cups). Emergence over time was analyzed using binomial generalized estimating equations (GEE) with a logit link, implemented in R for statistical computing using the geepack package. Fixed effects included time, seedpod design, and their interaction (weeks × seedpod design). Repeated daily observations were clustered by trajectory (specimen cup × seedpod design). Independence and first-order autoregressive (AR(1)) working correlation structures were evaluated, and the simpler structure was selected based on quasi-likelihood under the independence model criterion (QIC). Overdispersion was assessed using the performance package. Overall effects of seedpod design and time × design interactions were evaluated using Wald-type hypothesis tests. Post-hoc comparisons were conducted using estimated marginal means on the response scale with multiplicity-adjusted contrasts (Tukey or Holm adjustment). Where contrasts were reported as odds ratios, estimates and confidence intervals were computed on the log-odds scale and back-transformed. For the Northeastern experiment, formal germination inference focused on species exhibiting sufficient seedling emergence (*Betula lenta* and *Pinus strobus*), whereas species with negligible germination (*Acer rubrum* and *Tsuga canadensis*) were summarized descriptively.

**Supplementary Note 8. Emergence dynamics across seedpod designs in the indoor test**

For *B. lenta* and *P. strobus*, the GEE model indicated a steep increase in germination beginning around weeks 1–2, with wooden seedpods, particularly wooden seedpods (1 h), ranking highest across weeks 0–3. By week 3, germination reached ~62–64% for wooden seedpods (1 h) and ~60% for wooden seedpods (48 h), whereas the direct-sown control remained lower (~26–27%). Although most pairwise contrasts were uncertain after multiplicity correction, early comparisons in *P. strobus* indicated that wooden seedpods (1 h) exceeded the Control at week 1 (Holm-adjusted $p = 0.025$).

**Supplementary Note 9. Soil conditions in the Berkeley Forest.**

The soil is a highly organic material, predominantly behaving as an organic silt (OL) in terms of engineering properties. Although particle-size distribution shows a significant sand fraction with $D10 \approx 0.059$ mm, $D30 \approx 0.164$ mm, and $D60 \approx 0.546$ mm, these parameters are secondary because the high organic content dominates the soil's mechanical and hydraulic behavior. Atterberg limits indicate a liquid limit of 46% and a plastic limit of 35%, giving a plasticity index of 11%, which reflects low to moderate plasticity, largely controlled by the organic fraction. The soil at the site has a natural water content of 43%, which lies within the plastic range of the fines, indicating the soil is in a soft and deformable state. It exhibits high compressibility, low shear strength (Range: 10 - 200 kPa, depending on the compaction effort), and pronounced sensitivity to moisture changes. Hydraulic conductivity is expected to be significantly lower than would be predicted by conventional sand-based correlations, due to the presence of organic matter. Overall, the soil should be considered an organic silt, with properties controlled primarily by organic content rather than the mineral sand fraction, resulting in highly deformable and moisture-sensitive behavior.

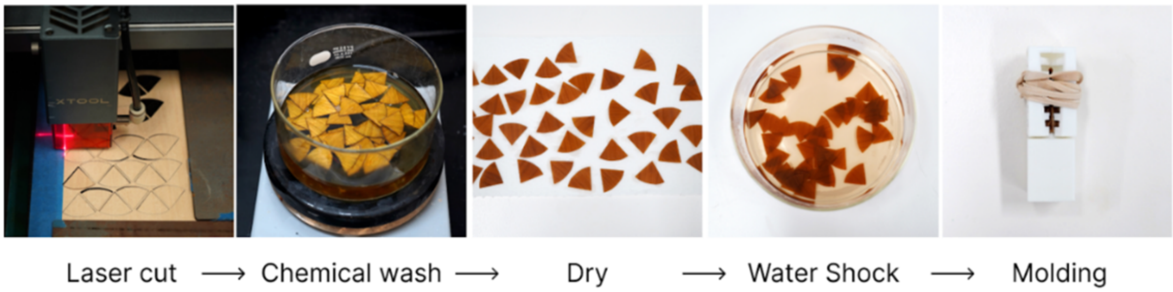


**Supplementary Figure 3. Schematics of the preparation of the wooden seedpod fabrication process.**

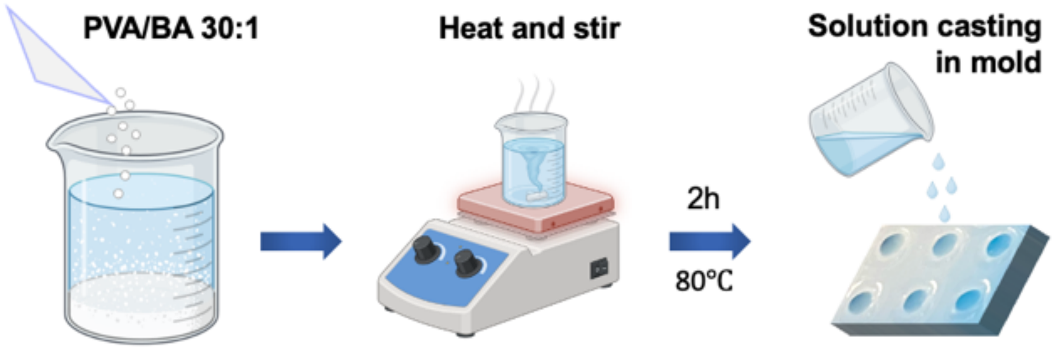


**Supplementary Figure 4. Schematics of the preparation of the poly(vinyl alcohol)/boric acid (PVA/BA) solution.**

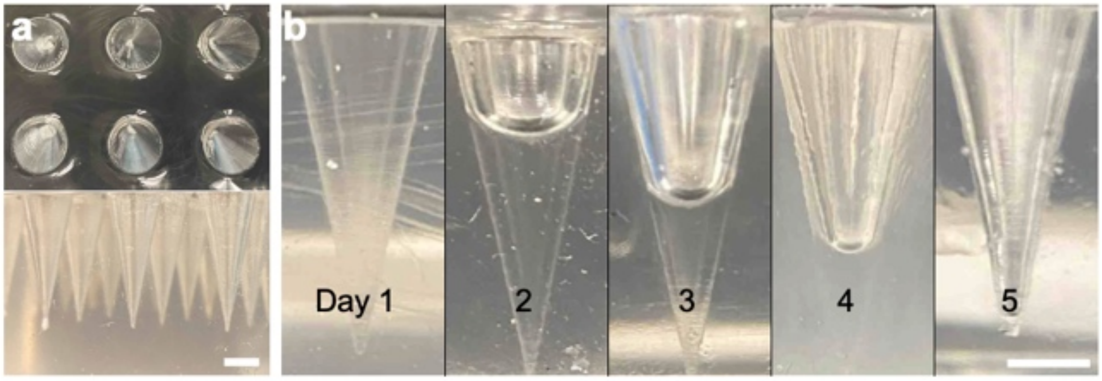


**Supplementary Figure 5. Evaporation of the PVA/BA solution in the mold. a.** Top and side views of the PDMS mold. **b.** Evaporation process of the solution in the mold. Scale bar, 5 mm.

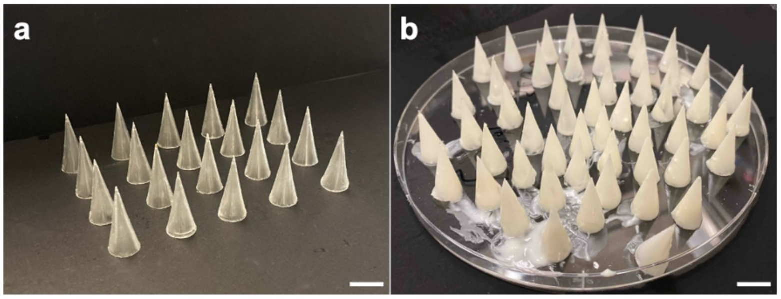


**Supplementary Figure 6. Experimental PVA/BA seedpods before (a) and after (b) wax coating.** Scale bar, 10 mm.

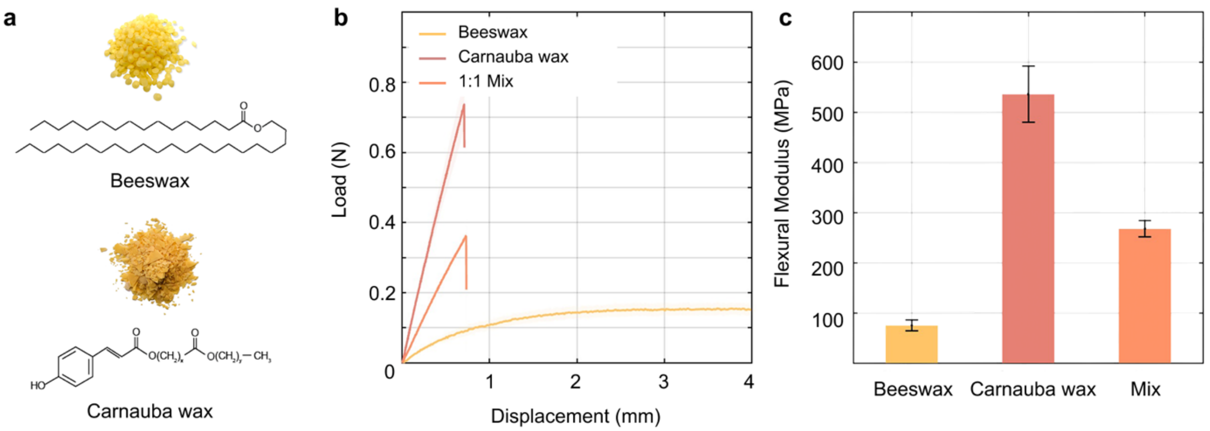


**Supplementary Figure 7. Mechanical characterization of beeswax, carnauba wax, and their 1:1 mixture. a.** Optical images of beeswax and carnauba wax, and their corresponding chemical structures. **b.** Three-point bending load-displacement curves for pure beeswax, pure carnauba wax, and their 1:1 mixture**. c.** Flexural modulus of beeswax, carnauba wax, and their 1:1 mixture (mean ± s.d., N = 5). Beeswax (75.1 ± 11.1 MPa), carnauba wax is very stiff (536.5 ± 56.1 MPa), and the mixture (268.1 ± 16.4 MPa), capturing both the rigidity of carnauba and the compliance of beeswax to yield a tunable delay-timer layer.

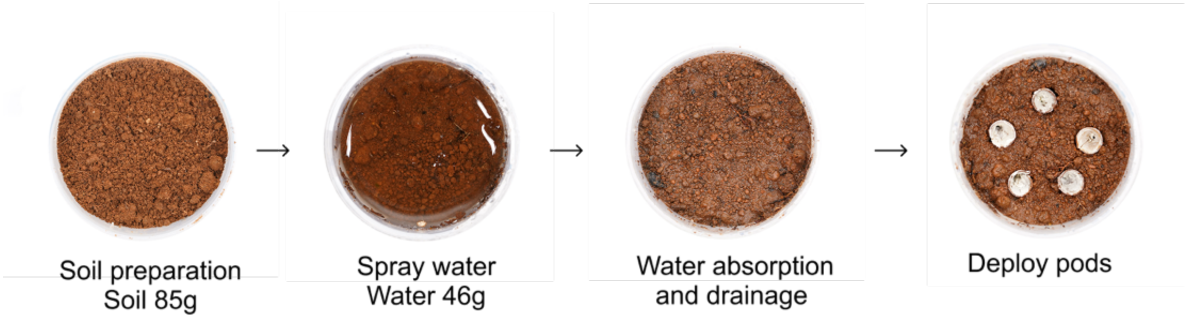


**Supplementary Figure 8 | Soil preparation for the indoor testing.** To investigate the effect of wax coating condition on seedpod opening time, we established a controlled test environment with a gravimetric water content of 43-46%. Water was applied by fine-mist spraying to ensure uniform wetting and minimize soil disturbance. Following a 20-minute drainage period to eliminate excess water, the engineered seedpods were arranged in a pentagonal configuration on the soil surface. The experiment was performed under controlled ambient room temperature (22 °C) and air humidity conditions (50%).

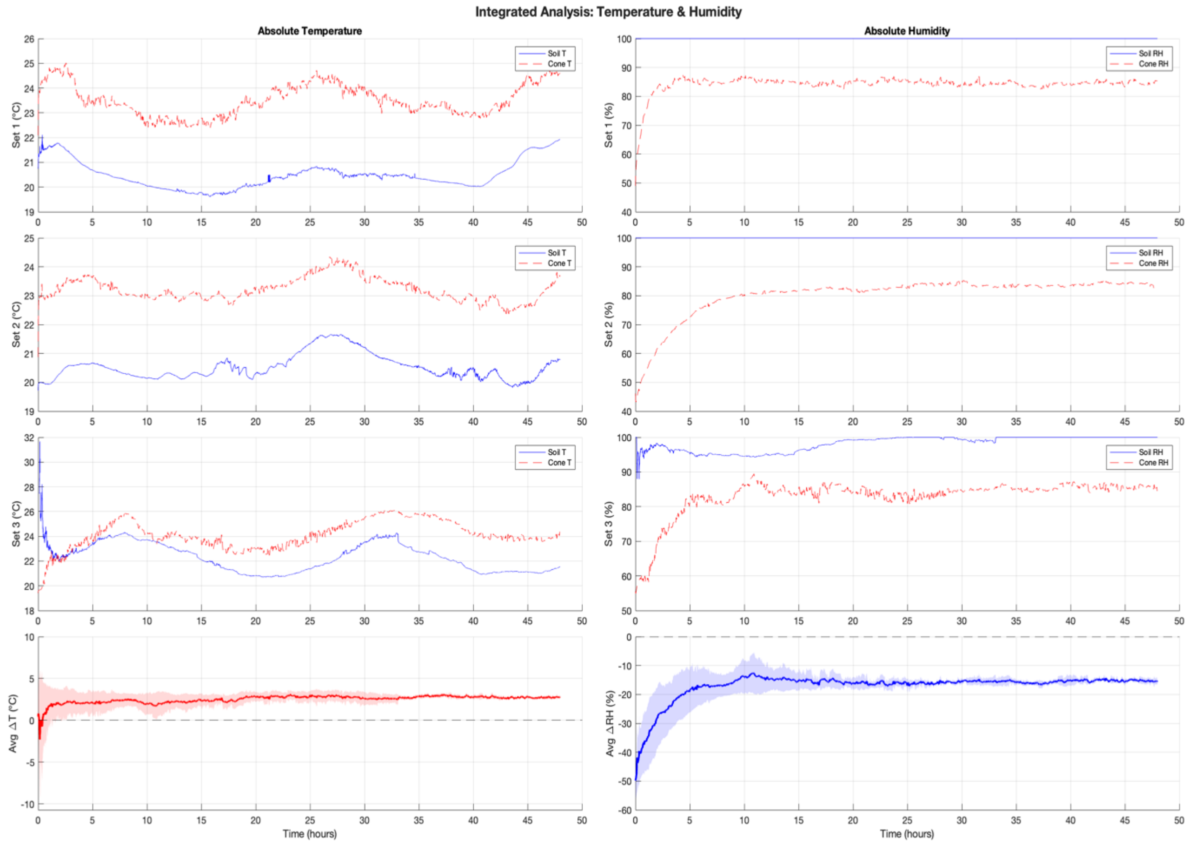


**Supplementary Figure 9. Monitoring temperature and humidity in soil and the wooden seedpod (1 h) over 48 hours.** Three sets of samples were monitored under indoor conditions in early spring in California (n = 3). Absolute temperature and humidity values are shown above, while the differences between the seedpod interior and the surrounding soil are presented below.

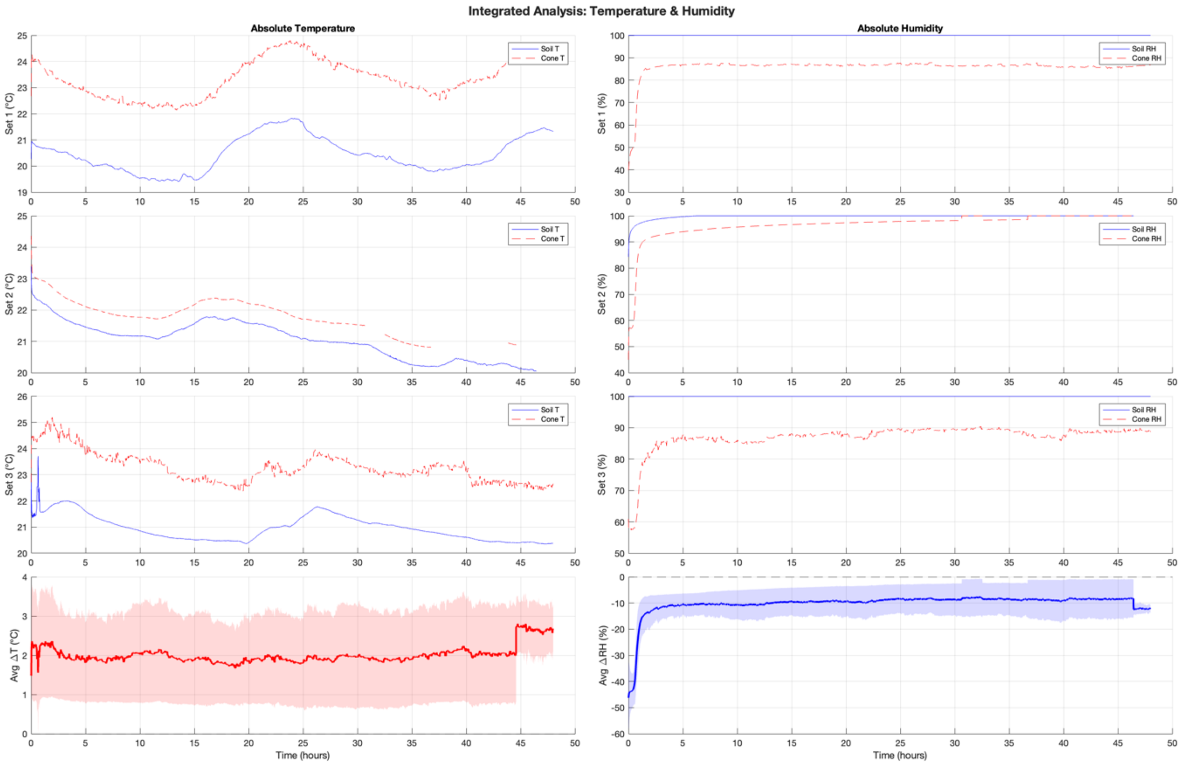


**Supplementary Figure 10. Monitoring temperature and humidity in soil and the PVA/BA seedpods (1 h) over 48 hours.** Three sets of samples were monitored under indoor conditions in early spring in California (n = 3). Absolute temperature and humidity values are shown above, while the differences between the seedpod interior and the surrounding soil are presented below.

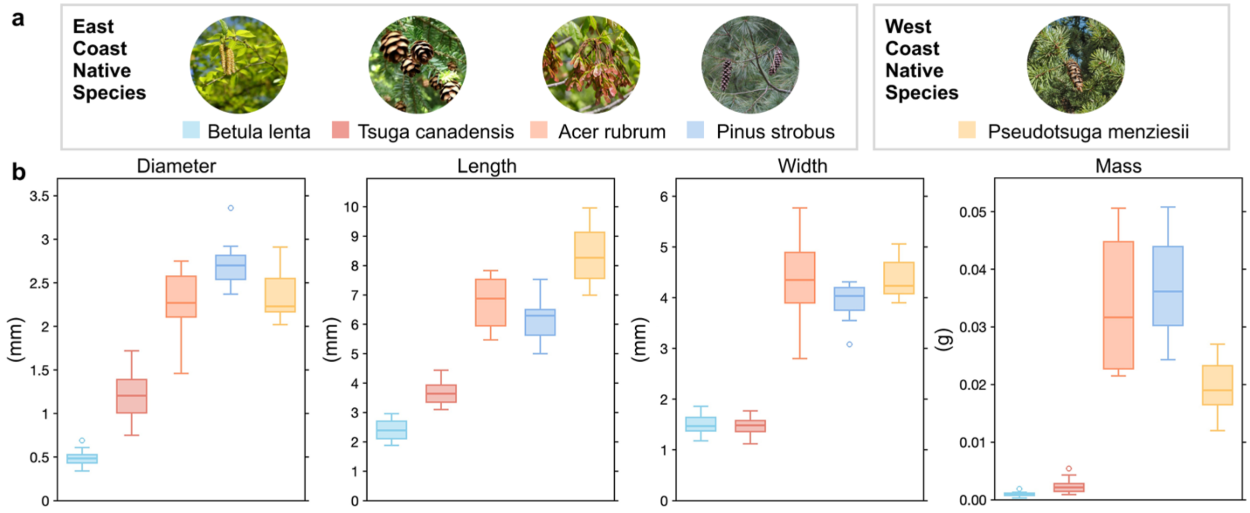


**Supplementary Figure 11. Seed morphology across species, including diameter, length, width, and mass, shown as box plots. a.** Field study plot with artificial irrigation and fencing system. **b.** Randomized plot design for each treatment to minimize location-specific effects. Data mean ± s.d., n = 40 per species.

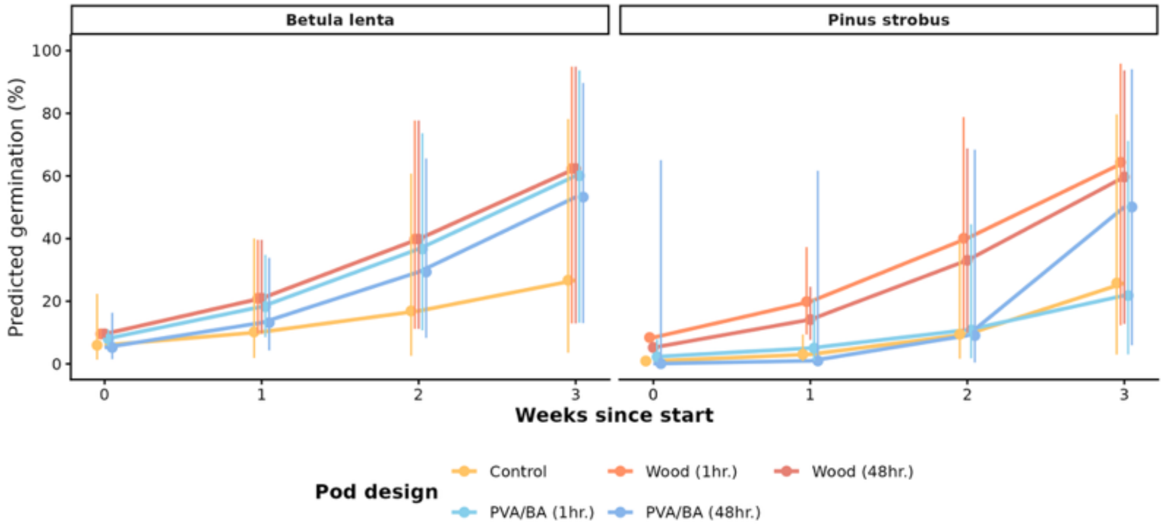


**Supplementary Figure 12. Germination (%) by seedpod design for *Betula lenta* and *Pinus strobus* from binomial GEE models with an AR (1) working correlation.** Points and lines show estimated marginal mean germination probabilities (back-transformed to %) at weeks 0–3 since deployment, and error bars indicate 95% confidence intervals. Facets separate species. Replication was n = 3 specimen cups per species × seedpod design (total n = 30 trajectories including Control), with repeated weekly observations clustered by trajectory.

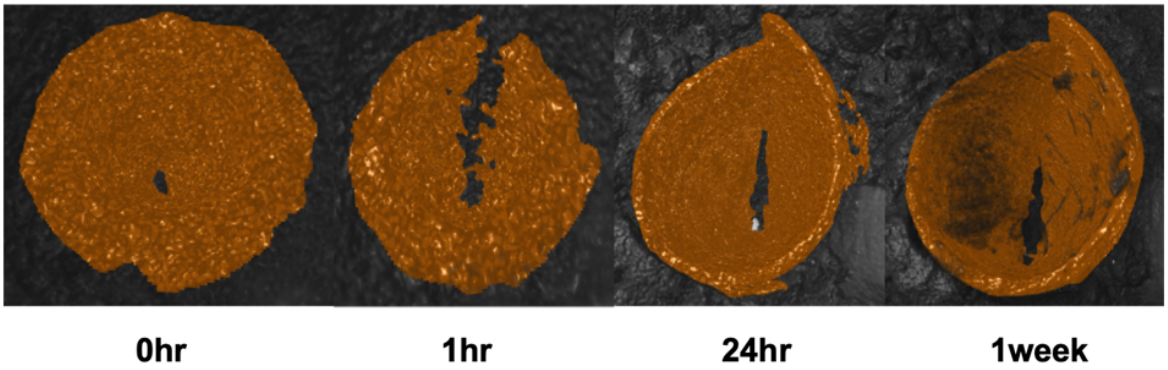


**Supplementary Figure 13. A wooden seedpod (1 h) crack-opening development through time, segmented after X-ray micro-computed tomography.**
